# High Throughput Viral Enumeration of Aquatic Ecosystems via Flow Cytometry

**DOI:** 10.1101/2024.10.21.619517

**Authors:** Madeline Bellanger, Pieter T Visscher, Richard Allen White

## Abstract

For the past 25 years, flow cytometry has been a gold standard for the direct measurement of viral-like particles (VLP) in aquatic ecosystems. Flow cytometry allows for higher throughput and costs less than alternative enumeration methods, leading to its broad usage in aquatic viral ecology. A major challenge associated with flow cytometry is the degradation of VLPs over time, making the use of high throughput plates not possible, thus lowering overall throughput. It has also been difficult to maintain a method with low contamination, high signal-to-noise ratios, and observations of real VLPs vs. fake particles. For these reasons, the use of flow cytometry has rapidly declined over the years due to the advent of massively parallel sequencing. Here, we describe a high throughput method in a 96 well plate format that provides a hands-free approach to viral enumeration. Our approach limits fake particles, noise levels, and cross-sample contamination. In a standard run, 60 samples can be measured for VLPs within 2.25 hours, which is ∼1 hour faster than the standard single tube approach and ∼1.5 hours faster than epifluorescence microscopy (EFM). Direct measurement of VLPs still provides a window into the viral-host interactions within aquatic ecosystems, which can be rapidly measured and resolved in a high throughput manner.

## 1. Introduction

The direct observation and measurement of viruses (i.e., as viral-like particles, VLPs) in aquatic ecosystems occurred 35 years ago, resulting in hundreds of publications using epifluorescence microscopy (EFM) or flow cytometry (FCM) to directly enumerate viruses (Bergh et al. 1989; Brussaard et al. 2000; 2004; 2010; Carreria et al. 2015; Noble et al. 1998; Patel et al. 2007). These measurements revealed that viruses are highly abundant across ecosystems globally, from the marine water column (Noble et al. 1998), marine sediments (Bergh et al. 1989; Danovaro et al. 2001), hot spring environments (Rachel et al. 2002), freshwater systems (Duhamel et al. 2006), soils (Williamson et al. 2003), microbial mats (Carreria et al. 2015), microbialites (Bellanger et al. 2024), and even the air (Prussin et al. 2015). Their cosmopolitan and ubiquitous nature on Earth has resulted in global estimates of 10^31^ VLPs, which accounts mainly for double-stranded DNA (dsDNA) phage abundances, and is currently only one order of magnitude greater than the estimated bacterial abundances (Hendrix et al. 1999; Mushegian, 2020). This may still be an underestimate due to the lack of direct methods to measure and decipher all nucleic acid types individually within a complex VLP mixture (e.g., single stranded DNA vs. dsDNA, or ssRNA vs. dsRNA, or RNA vs. DNA) (White III et al. 2021; White III, 2021). Giant viruses may also be filtered out and thus missed by many measurements in aquatic ecosystems due to the use of 0.22 μm filters (Philippe et al. 2013; Sun et al. 2014; White III et al. 2021; White III, 2021). Viruses have been estimated to be 10^30^ within the global oceans alone and with global biomass estimates, suggesting an underestimation with further methods needed to enumerate viruses on a global scale (Suttle, 2007; Bar-On et al., 2018).

Viruses have been found to be global players in nutrient cycling and transport, ecosystem services, global biogeochemical cycling (e.g., carbon and nitrogen pool), marine carbon pump (i.e., the viral shunt), horizontal gene transfer, and food web dynamics (Wilhelm & Suttle, 1999; Fuhrman, 1999; Weinbauer, 2004; Suttle, 2007; Rohwer et al. 2009; Jiao & Zheng, 2011; Breitbart, 2012; Lara et al. 2017; Irwin et al. 2022; Wang et al. 2022; Carreira et al. 2024). Yet even with all their critical ecological roles tools for direct measurement of VLPs have been limited to low throughput methodologies such as single tube FCM or single EFM. A method that allows for high throughout direct measurement of VLPs is needed to provide robust and accurate global estimations of the total VLPs present on Earth that can scale to such global sampling. Enumeration via direct counting is the first path towards understanding the viral role within ecosystems and therefore must be further examined and developed.

EFM and FCM are the most broadly applied tools for enumeration of VLPs within ecosystems (Noble & Fuhrman, 1998; Marie et al. 1999; Brussaard, 2004). EFM has been used less due to the high cost of Anodisc filters (>$15 a slide), which are often difficult to obtain, and this approach is thus not amenable to high throughput (Bellanger et al. 2024). Wet mount methods by comparison are cheaper around $0.18-0.84 per slide (Bellanger et al. 2024). However, EFM always results in lower throughput than FCM, even with the current single tube non-plate based approach we discuss here. EFM is more labor intensive to obtain counts, which include individual slide focusing, capturing many fields within the slides (∼4 or more fields), images are currently counted by eye, and lower throughput when compared to FCM. However, a new software has been shown using computer vision to automate the counting of VLPs within EFM slides (Figueroa III et al. 2024). Issues outlined above besides, FCM allows for sensitive detection, quantification, and rapid analysis of viral populations in high throughput (Brussaard et al. 2010). Major advancements in FCM have occurred over the last decade including sensitive detection of nanoparticles at the 40 nm range with nano-scale FCM (Beckman, 2024).

EFM and FCM have been criticized for measuring ’fake particles’ instead of VLPs (Forterre et al., 2013). These fake particles that are commonly found in FCM and EFM methods include membrane-derived vesicles (MDVs), free extracellular DNA (FED), gene transfer agents (GTA), and cell debris (Forterre et al., 2013). Other potential contaminants include ribosomes, which can be co-purified with viruses and are difficult/complicated to remove (Conceição-Neto et al., 2015). The method recently described by Bellanger et al. (2023) improved previous methods by adding a chloroform step, which reduces contamination by MDVs, and ribosomes. Bellanger et al. (2023) also added a benzonase nuclease step, which removed FED due to lack of protection for free DNA. Most enveloped viruses are sensitive to chloroform due to loss of stability as a result of/resulting from envelope removal (e.g., coronaviruses and mimiviruses) (Conceição-Neto et al., 2015). Some non-enveloped also can be sensitive to chloroform (e.g., *Inoviridae*) (Conceição-Neto et al., 2015). This, however, is a trade-off as removing the MDVs and ribosomes may be needed for higher quality results.

Here, we present a high throughput method of FCM to enumerate viruses in various types of aquatic environments (freshwater, hypersaline, and marine), ensuring sample stability. Our method utilizes the well plate loader on a Beckman Coulter CytoFlex (Beckman Coulter, Brea, CA, USA), building on protocols developed by Brussaard et al. (2010), incorporating the suggestions of Forterre et al. (2013), and ensuring all samples are stable throughout the entire run. Enumeration of viruses within aquatic ecosystems could illuminate the virosphere, its interactions and mechanisms on a global level.

## 2. Materials and Methods

### Sample Collection

Freshwater, hypersaline, and coastal marine samples of ≥1 L were collected. Green Lake meromictic freshwater samples were collected from the mixolimnion in October 2023 (FGL, Green Lakes State Park, New York, 43.049°N, 75.973°W). Great Salt Lake hypersaline water was collected in July 2020 (GSL, Antelope Island State Park, Utah, 41°N, 112°W, near Layton, Utah). Coastal marine samples from Wrightsville Beach were collected in December 2023 (WBS, National Data Buoy Center Station 41038, 34.141° N, 77.715° W, near Wilmington, North Carolina). An previously isolated cyanophage from a Green Lake microbialite was used as a positive control during initial testing of the method.

### Preparation of Nucleic Acid Stain Working Stock

The working stock was created from a commercial stock of SYBR Gold (Invitrogen S11494), starting with thawing in the dark at room temperature. Once thawed, the commercial stock was vortexed for 10 s at medium-high speed (∼1,000 RPM), then centrifuged at 2,000 RCF in a microcentrifuge for five minutes. The commercial stock was then diluted 1:200 with autoclaved and filtered (0.22 μm PVDF Millipore GVWP06225) molecular biology grade water. The working stock was filtered (0.22 μm PVDF filters) before small volumes (∼300 μL) were aliquoted into black microcentrifuge tubes and stored at -20°C until use. Multiple freeze-thaw cycles of the working stock should be minimized.

### Preparation of TE Buffer Working Stock

The TE buffer working stock was created by diluting 10X Tris-EDTA with molecular biology grade water to achieve a 1X solution. The solution was immediately autoclaved on a liquid 15 setting. Before each use, a small amount (≤50 mL) of prepared TE buffer working stock was aliquoted into a new tube and filtered down to 0.2 µM using sterile syringe filters.

### Well Plate Planning

To achieve high throughput, a 96 well plate was used. Each well plate was planned out ahead of time. To avoid over-counting of events and to maintain the health of the flow cytometer, a well containing a 10% bleach solution was ran for every five sample wells to clean the flow cytometer (**Supplemental Figure 1**). If bleach is unavailable, 70% ethanol or a specialized flow cytometer cleaning solution may be used. Additionally, a well with nuclease free water was ran after each bleach well (**Supplemental Figure 1**). This well serves as both a way to rinse any residual bleach and a way to keep track of the noise levels of the machine. This blank could also be the TE buffer working stock. Running blanks consistently ensures that the noise levels of the cytometer can be monitored and stay low. Ultrafiltrate from the same location as the sample was used to determine the base noise level of the sample. An ultrafiltrate blank was ran for every five sample wells after the nuclease free water well to ensure noise levels of the sample stay low throughout the analysis (**Supplemental Figure 1**). The ultrafiltrate blank was prepared the same as the sample.

The amount of sample needed to be fixed was calculated based on the dilution factor. Here, 11 wells were reserved for bleach, 12 wells were reserved for blanks, one for TE buffer, and 12 for ultrafiltrate. This left 60 wells for the sample (**Supplemental Figure 1**). Within the 96 well plate, the volume used in each well was 150 µL. A total volume of 9 mL of diluted sample was needed, but was rounded up to 10 mL to allow for any errors and easier calculations. The dilution factor will be dependent on the amount of events measured in one minute of acquisition (see *Flow Cytometry Analysis* for optimal event rate). A 1:100 dilution was used in this study, resulting in 100 µL of environmental sample needed for a single well plate. Similarly, a total volume of 1.8 mL of diluted ultrafiltrate was needed, but was rounded up to 2 mL. Using the same dilution factor as the sample (100X), 20 µL of ultrafiltrate was needed for one well plate.

### Preparation of Samples

An overview of the sample preparation can be seen in **Figure 1**. As detailed in Bellanger et al. (2023), the samples were filtered through 0.22 μm PVDF filters and concentrated using Centricon-70 plus centrifuge filters (Millipore UFC703008, 30 kDa). A portion of the samples were chloroform treated to observe the benefits obtained from chloroform treatment. The calculated amount of sample above (100 µL) was aliquoted into a low binding 1.5 mL tube. Samples were then fixed according to Bellanger et al. (2023) at a final concentration of 0.5% glutaraldehyde.

**Figure 1.**
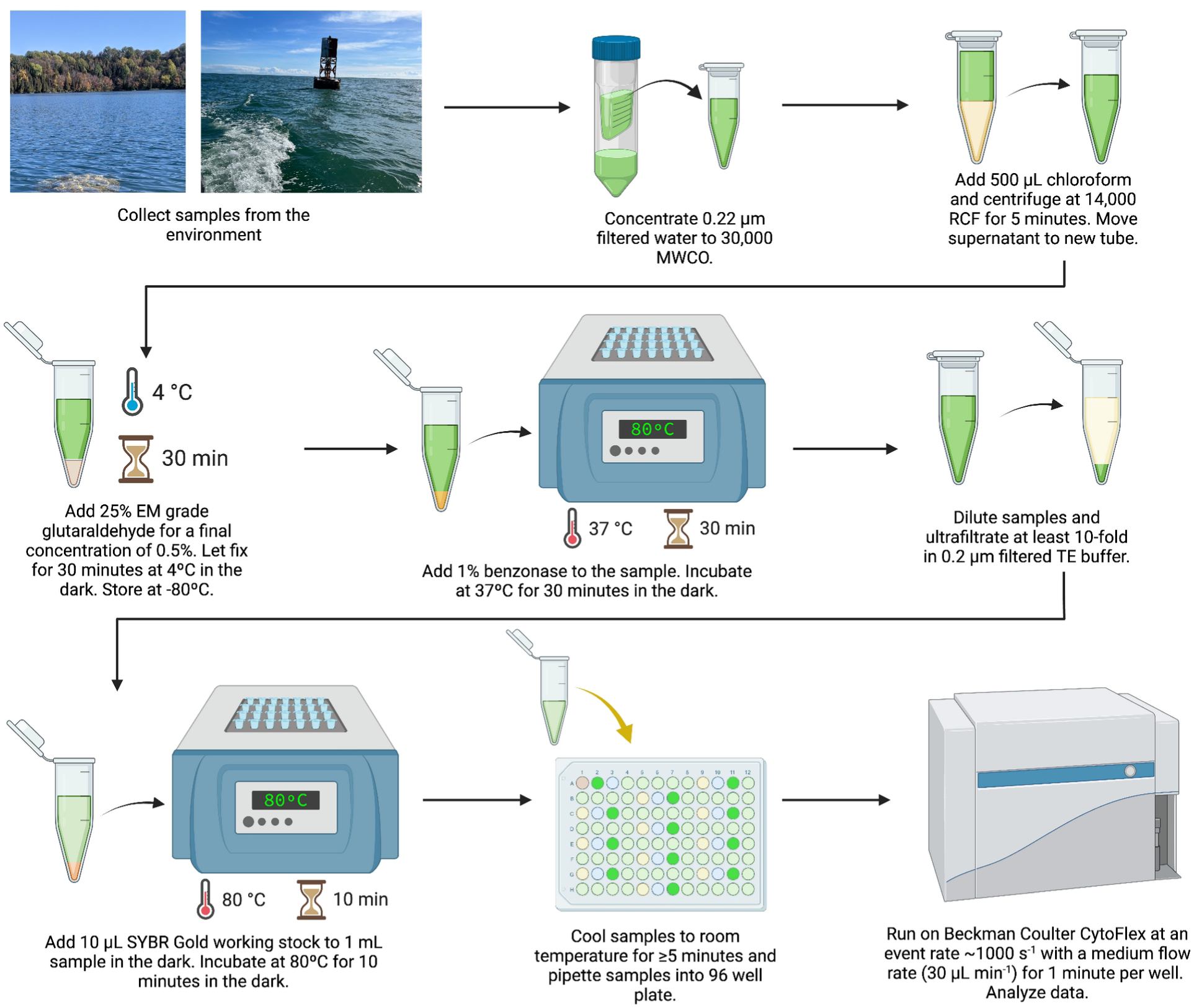
Flow diagram of flow cytometry protocol. This diagram provides a detailed methodological walk-through of the preparation of water samples for VLP enumeration. The entire protocol, including running the entire 96 well plate on the flow cytometer, takes ∼3 hours.

Fresh low binding 1.5 mL tubes were prepared, so that 1 mL of the final diluted sample was in each tube, in addition to two tubes for ultrafiltrate. Heat blocks were preheated to 37°C and 80°C. The SYBR Gold working stock and, when applicable, the samples were thawed in the dark to room temperature. To ensure free floating nucleotides were not being counted as VLPs, a portion of the samples were treated with benzonase for a final concentration of 1% (250 U) and incubated at 37°C for 30 minutes. Samples not treated with benzonase were thawed and immediately diluted.

To prepare the samples for dilution, 10 µL of the sample was aliquoted into each of the 1.5 mL sample tubes. To achieve a 100-fold dilution, 990 µL of filtered TE buffer working stock was added to the samples and pipetted to mix. Samples should be diluted at least 10-fold for an optimal event rate. If the contents of a sample and the potential dilution factor were unknown, the undiluted sample was ran on the flow cytometer using tube loading to get an initial event rate. Multiple dilution factors were then tested to figure out the most optimal dilution. Dilution factors can be as high as 1,000X for high-titer lysates (Bonar & Tilton, 2017). The ultrafiltrate blanks were prepared in the same manner as the samples. The rest of the steps were performed on both the samples and the ultrafiltrate blanks (hereby referred to as "samples").

The samples were dyed with 10 µL of the SYBR Gold working stock and incubated at 80°C for 10 minutes in the dark. After incubation, the samples were cooled to room temperature for at least 5 minutes while being pipetted into the well plate. All well plates were then ran on a Beckman Coulter CytoFlex using well plate loading and a violet side scatter laser configuration to observe nanoparticles. An entire well plate takes roughly 2.25 hours to run when using the Auto Record function.

### Flow Cytometry Analysis

Each well was ran to obtain an optimal event rate between 100 and 500 events s^-1^ using a medium flow rate of 30 µL min^-1^ for one minute of acquisition. The TE buffer working stock well was ran first to figure out the base noise level of the cytometer. The nuclease free water was then ran to cleanse the cytometer. The ultrafiltrate blank well was then ran to find the base noise level of the sample. Low coincidence and background noise was detected, with ∼200 events s^-1^ in one minute of acquisition at a medium flow rate and little to no events in the area of interest. The acquisition settings may be adjusted at this point to lower noise levels. The gain acquisition settings for all experiments were as follows: FSC 3000, SSC 1680, V-SSC 1, FITC 480, PE 200, KO525 45. The threshold was set to V-SSC at 3000. If the event rate is too high (i.e., >500 events s^-1^), VLP populations will not be discernable and samples will need to be diluted further. Alternatively, if the event rate is low (i.e., <100 events s^-1^), more noise will be detected and the samples will need to be diluted less. If a flow cytometer that is not equipped with the violet side scatter laser configuration is used, the area of interest will be in a different quadrant than that shown in **Supplemental Figure 2**, as resolution of the cytometer will not be as high.

The following histograms were used for analysis: FSC-A, SSC-A, Violet SSC-A, FITC-A. Additionally, the following dot plots (X axis vs Y axis) were used for analysis: FSC-A vs SSC-A, FSC-A vs Violet SSC-A, FITC-A vs SSC-A, FITC-A vs Violet SSC-A.

Gates should be set for each sample that include VLPs and exclude the background noise. A gate for each viral population should be put in place (**Supplemental Figure 2**). A gate for noise can be drawn on the ultrafiltrate plots and the nuclease free water plots to ensure that those areas are not included in VLP gate(s). The amount of events in the VLP gate(s) will be used to find the concentration of VLPs in the sample. The VLP concentration can be calculated using the formula developed by Maltseva & Langlois (2022). Analyses in this experiment were conducted using the CytExpert software (Beckman Coulter, Brea, CA, USA), however, further analyses can be performed using softwares like FCS Express (De Novo Software) and FCMPass (Welch et al., 2020).

### Statistical Analysis

All statistical analyses were performed in R. For experiments with a CV higher than 15%, a statistical test was conducted to identify degradation. For those experiments, the data for each well in a row was gathered as a single population. Normality was determined for each population, and, if applicable, variance was compared to row A (**Supplemental Tables 1-3**). A statistical comparison (i.e., Student’s T test, Welch’s T test, or Wilcoxon Rank Sum) was conducted between row A and each subsequent row following to determine if the sample was stable throughout the entire experiment (**Supplemental Tables 1-3**). For samples where degradation was detected, wells were grouped into three with a dynamic sliding window approach. Each group was compared to the first group of three to find the exact point of degradation (**Supplemental Table 3**).

### Data and Code Availability

The flow cytometry data and code used in this work are publicly available on GitHub (https://github.com/raw-lab/HTSflow) and OSF (https://osf.io/ade4b/). The full protocol can be found on <u>protocols.io</u>.

## 3. Results

### Untreated Samples

The viral abundance in samples from FGL averaged 1.82 x 10^7^ VLPs mL^-1^, with a CV of 29.1% (**Figure 2, Supplemental Table 4**). Over the course of the well plate run, no significant decline in VLP concentration was observed (**Supplemental Table 1, p > 0.05**). Samples from GSL showed similar results, with viral abundances averaging 1.16 x 10^7^ VLPs mL^-1^, with a CV of 36.82% (**Figure 2, Supplemental Table 4**). Samples in the GSL well plate also showed no significant decline in VLP concentration (**Supplemental Table 2, p > 0.05**).

**Figure 2.**
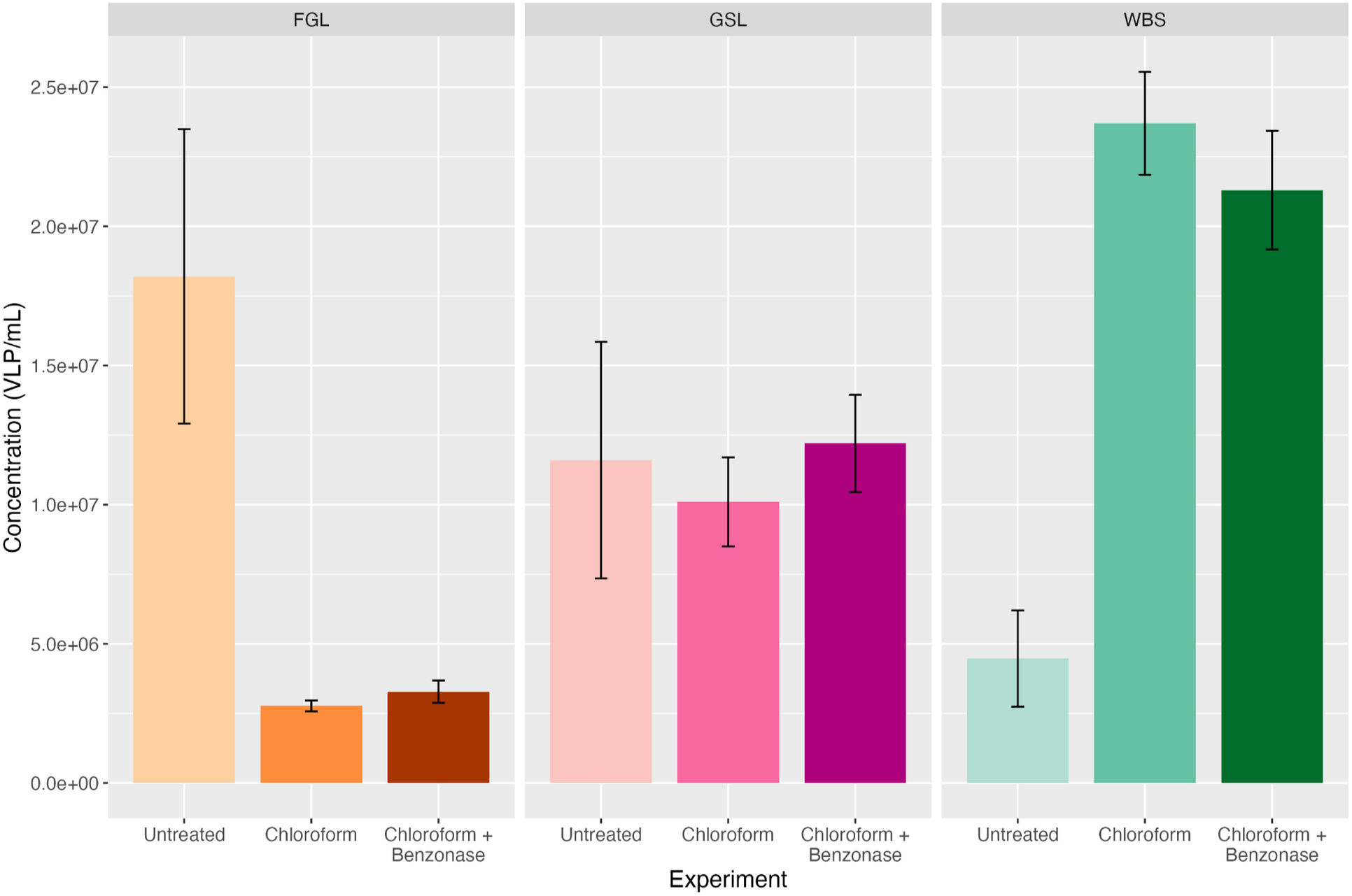
Average VLP mL^-1^ for FGL, GSL, and WBS using different sample preparation methods. Error bars represent standard deviation values.

Samples from WBS had an average viral abundance of 4.47 x 10^6^ VLPs mL^-1^, with a CV of 38.74% (**Figure 2, Supplemental Table 4**). Contrary to FGL and GSL, WBS showed a significant decline beginning at row F, roughly 1 hour and 20 minutes into the run (**Supplemental Table 3, Supplemental Figure 3, p < 0.01**). Counts remained briefly constant before declining in row G, roughly 1 hour and 35 minutes into the run (**Supplemental Table 3, Supplemental Figure 2, p < 0.01**).

### Chloroform Treated Samples

The viral abundances of FGL samples that were treated with chloroform averaged 2.77 x 10^6^ VLPs mL^-1^, which was lower than the untreated samples (**Figure 2, Supplemental Table 4**). The CV was 7.01%, so no statistical test for degradation was conducted (**Supplemental Table 4**). Chloroform-treated GSL samples had higher viral abundances than those from FGL, averaging 1.01 x 10^7^ VLPs mL^-1^, nearly equal to the untreated GSL samples (**Figure 2, Supplemental Table 4**). A statistical test was not conducted due to the CV being under 15% (**Supplemental Table 4**). The treated WBS samples had an average viral abundance of 2.37 x 10^7^ VLPs mL^-1^, higher than the untreated samples (**Figure 2, Supplemental Table 4**). This is likely due to fewer large particles, allowing the cytometer to identify smaller particles and VLPs as events. The CV was low, at 7.82%, so no statistical test for degradation was conducted (**Supplemental Table 4**).

### Benzonase Treated Samples

Samples were also treated with both benzonase and chloroform. These FGL samples had viral abundances that averaged 3.28 x 10^6^ VLPs mL^-1^ (**Figure 2, Supplemental Table 4**). This average was lower than the untreated FGL sample, but slightly higher than the chloroform-only treated FGL sample. The CV was 12.24%, so no statistical test for degradation was conducted (**Figure 3, Supplemental Figure 4, Supplemental Table 4**). The benzonase and chloroform treated GSL samples again had higher viral abundances than FGL, averaging 1.22 x 10^7^ VLPs mL^-1^ (**Figure 2, Supplemental Table 4**). This average was nearly equal to both the untreated samples and the chloroform treated samples. This sample had a CV of 14.41%, so a degradation test was not needed (**Figure 3, Supplemental Figure 4, Supplemental Table 4**). The benzonase and chloroform treated WBS samples had an average viral abundance of 2.13 x 10^7^ VLPs mL^-1^ (**Figure 2, Supplemental Table 4**). This average was higher than the untreated samples and nearly identical to the chloroform treated samples. Similar to the chloroform treated samples, the higher average is likely due to better distinction of VLPs, as fewer unwanted particles would be identified. The CV was low, at 10.03%, so no statistical test for degradation was conducted (**Figure 3, Supplemental Figure 4, Supplemental Table 4**).

**Figure 3.**
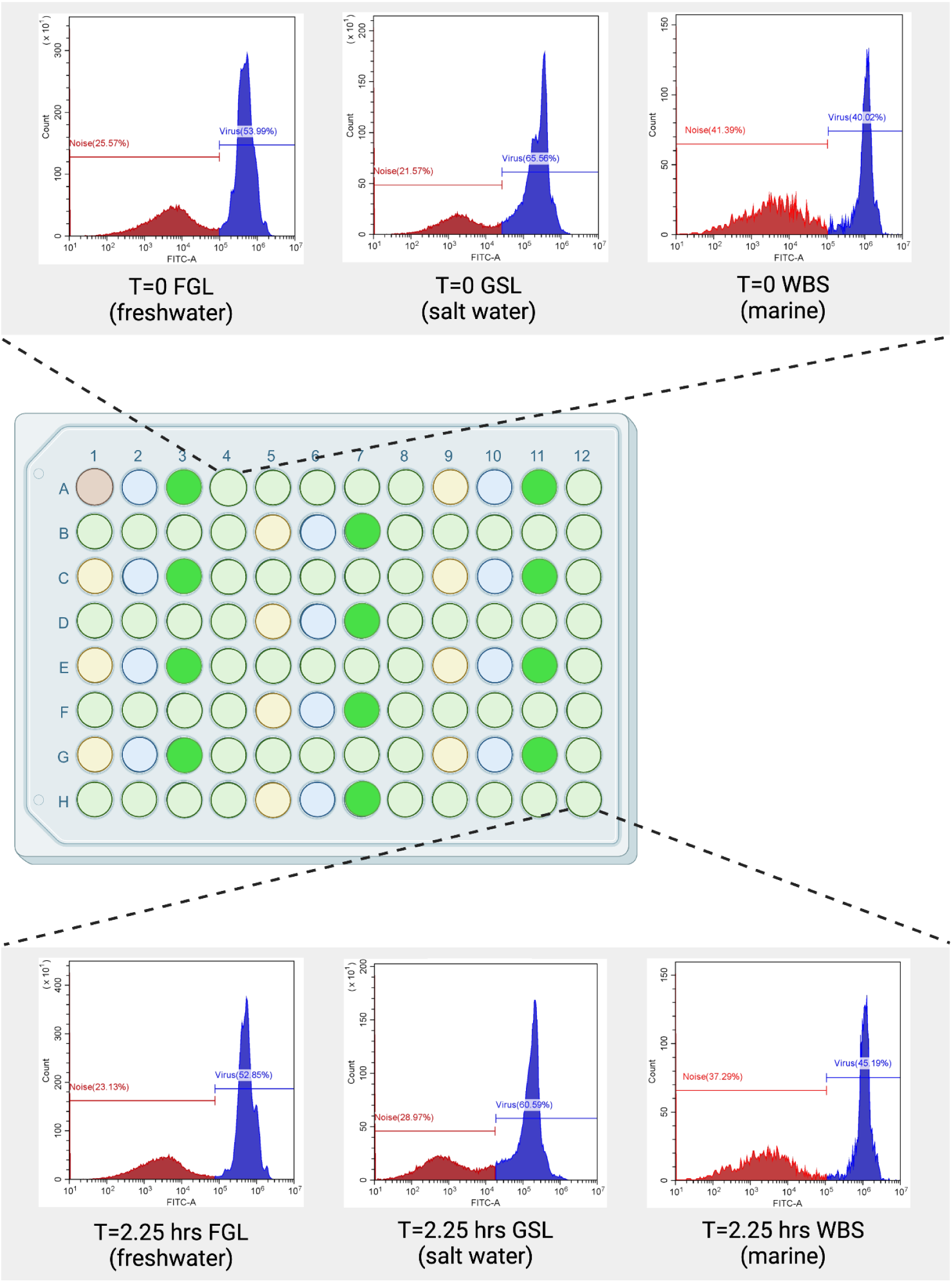
Time series results for each sample. Histogram plots show the level of FITC fluorescent emission intensity. Gates that are colored blue are areas that contain viruses. Gates that are colored red are noise areas. VLP concentrations stay consistent throughout the entire experiment over the course of ∼2.25 hours.

### Methodology Comparisons/Comparison of the Various Treatments

The untreated samples showed a large amount of variability throughout the entire run (**Supplemental Table 4**). The addition of chloroform to the samples greatly improved this variability, suggesting its beneficial use for sample stability (**Figure 2**). Additionally, using benzonase allowed for counts being attributed to viral particles, rather than extracellular DNA (**Figure 2**). The sample preparation used here, first designed for EFM by Bellanger et al. (2023), allows for a direct comparison of enumeration methods with the FGL and GSL samples (**Figure 4**).

**Figure 4.**
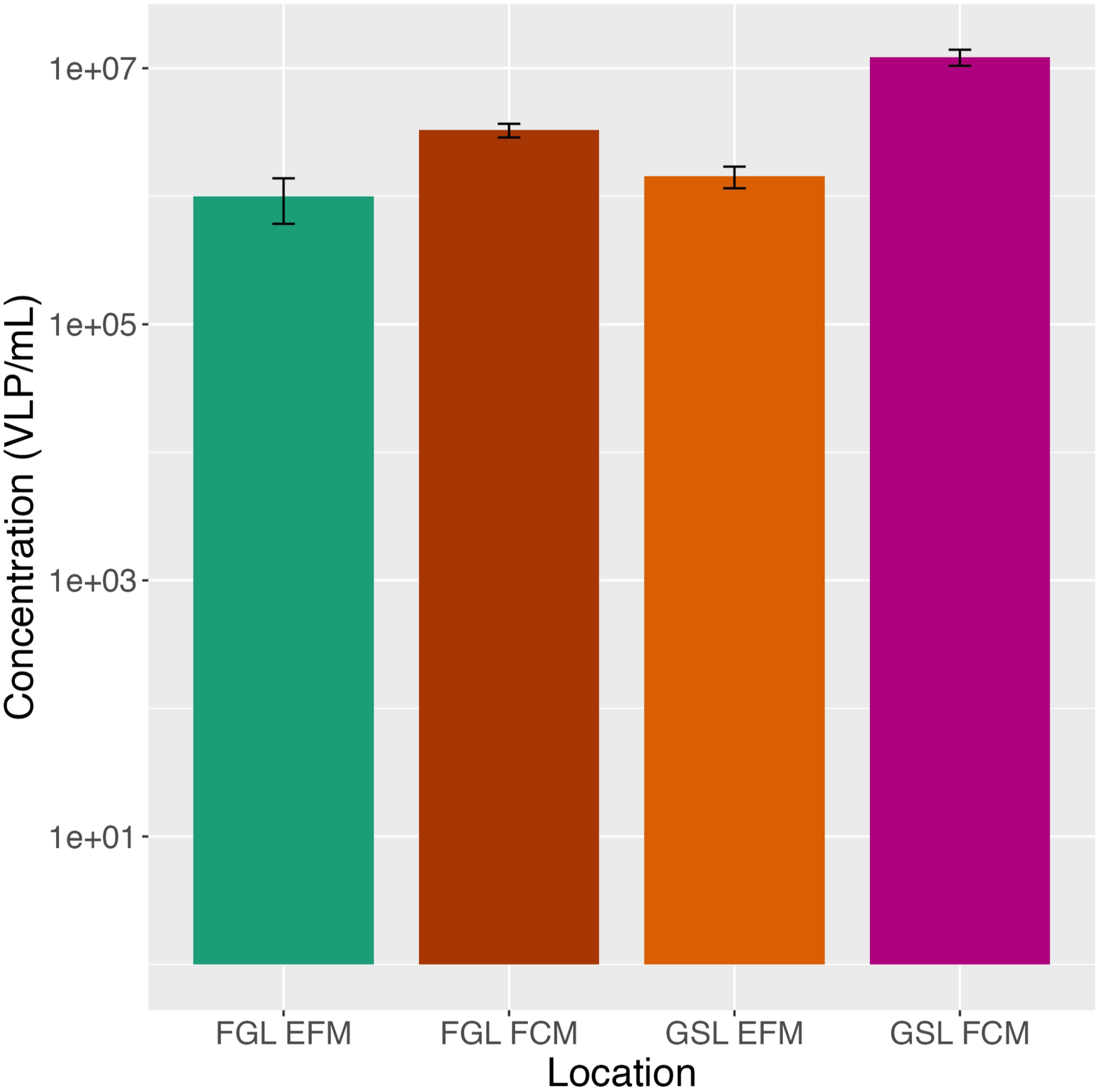
Average VLP mL^-1^ for FGL and GSL using EFM and FCM. EFM counts were obtained from Bellanger et al., 2023. Error bars represent standard deviation values.

The FGL samples had the lowest VLP concentrations of all locations (**Supplemental Figure 5, Supplemental Table 4**). The VLP concentrations of both the GSL and WBS samples were nearly an order of magnitude higher than the FGL samples (**Supplemental Figure 5, Supplemental Table 4**). Overall, these values were higher to those previously measured with EFM, likely due to a higher resolution through FCM (**Figure 4, Supplemental Table 4**).

Overall, our method provides the cheapest/least expensive and quickest/fastest way to enumerate VLPs in aquatic environments. A time and cost analysis is shown in **Table 1**. When counting an entire well plate, our FCM method is 25 times cheaper than using the EFM method described by Bellanger et al. (2023). Additionally, it is nearly an hour and a half quicker than EFM and nearly 30 minutes quicker than the traditional tube loading method per sample. When conducting EFM or flow cytometry using tube loading, the operator/technician has to be present for each measurement. However, our method uses the Auto Record function of the CytExpert software, allowing for the operator/technician to complete other tasks while the machine takes measurements.

**Table 1.** Time and Cost Analysis of Methods. Cost analysis of enumeration approaches for counting viruses in water. The cost of consumables (pipette tips, microcentrifuge tubes, etc.) is estimated to be the same for each method. While base EFM costs were obtained from Bellanger et al., 2023, costs were updated to reflect current prices at the time of calculation. A sample is denoted as one slide containing 10 µL of sample for EFM and 1 mL of 1:100 diluted sample for FCM.

| Method | Min Per Sample | Cost Per Sample (USD) | Min Per Well Plate | Cost Per Well Plate (USD) |
| --- | --- | --- | --- | --- |
| EFM (Bellanger et al., 2023) | 3 | \$0.38 | 216** | \$27.25** |
| FCM Tube Loading | 2 | \$0.10* | 192 | \$1.09* |
| FCM Well Plate Loading | 1.38 | \$0.10* | 132.5 | \$1.09* |

\*Prices calculated per mL of sample when diluted 1:100.
\*\*Calculated based on 72 samples for EFM, as that is what corresponds to the number of sample wells (60 sample wells + 12 ultrafiltrate wells) in a well plate.

## 4. Discussion

Our optimized FCM protocol presented here uses advances in technology that allow for better detection of viral populations over noise, while avoiding FEDs, MVDs, and cellular debris. Chloroform treatment showed improvements in sample stability, especially of marine samples. The combined chloroform and benzonase addition prevented VLP degradation and CVs decreased (**Figure 2**).

All samples analyzed had VLP abundances similar to those reported in various aquatic environments at ∼10^6^ to 10^7^ (Clasen et al., 2008). Overall, WBS had the highest VLP concentration, with 2.13 x 10^7^ VLPs mL^-1^ compared to GSL and FGL (**Supplemental Figure 5**). This is well within range of other coastal marine VLP abundances, even with the addition of chloroform and benzonase (Finke et al., 2017). This demonstrates that the treatments used in our method do not have a negative effect on the VLP concentrations but instead allow for improved sample stability.

Viral concentrations may vary seasonally and spatially with depth and relation to the shore (Brum et al., 2016). Seasonal measures of viral abundances are needed within aquatic ecosystems to enumerate any variation, especially in areas that experience severe droughts like GSL. Identifying the seasonality of viral abundances could allow for a better prediction of, e.g., the effects that global warming has on the microbial community. Additionally, observing deviations from normal seasonal variations may provide insight to the ecosystem health and interactions occurring within. Documenting these seasonal trends can allow for better understanding of how aquatic ecosystems respond to both natural disturbances and anthropogenic impacts.

Currently, our method enumerates viruses regardless of the nucleic acid type or strandedness and can account for large viruses. Giant viruses, including pandoraviruses, are commonly filtered out during standard filtration of 0.22 μm (Philippe et al., 2013). Chloroform without filtration has been useful in removing bacteria to isolate jumbo and megaphages (Saad et al., 2019). Antibiotics and fungicides may be useful for membrane-bound giant viruses like pandoraviruses to remove bacteria and fungi in enumeration studies (Philippe et al., 2013).

Further improvements are needed in fluorescent dyes to distinguish single-stranded vs. double-stranded nucleic acids when mixed (White III et al. 2020; White III, 2022). DAPI and Yo-Pro-I were previously used for viral enumeration, but were too dim compared to SYBR-based options (Patel et al., 2007). SYBR Green I and SYBR Gold showed equivalent counts for virus and bacterial enumeration (Patel et al., 2007). SYBR green I/II or SYBR gold can be used for RNA virus staining (Patel et al., 2007), but not in RNA and DNA virus mixed samples. Acridine orange may be selective for RNA over DNA in fluorescence shift and effective for ssDNA phages (Darzynkiewicz, 1990; Mayor & Hill, 1961). The development of FCM methods using various nucleic acid, protein, and lipid stains may help to distinguish intact viral particles and expose their presence, regardless of their nucleic acid type, strandedness, or size. Thus, further improvements of strains are needed to capture RNA viruses when mixed with DNA viruses.

The enhanced sensitivity and scalability of our method enables more comprehensive studies of viral ecology. The expansive use-cases of our method could allow for more insight into virus-host dynamics and viral roles in nutrient cycling. Additionally, our method allows for detection of viral pathogens in water systems, providing an efficient way to improve water quality management. Further improvements are needed to directly enumerate viruses within natural environments beyond what is presented here. Modified sample preparation paired with our method could allow for viral enumeration of a wider range of sample types outside of aquatic environments, like soil, microbialites, and exopolymeric substances, revealing the full extent of the world’s virome.

## Acknowledgements

We acknowledge the University Research Computing and the College of Computing and Informatics for computational and logistical support. We also acknowledge Bryan Fulghum, Scott Perry, and Ryan Perry for their assistance in sample collection.

