## Supplemental data for "High Throughput Viral Enumeration of Aquatic Ecosystems via Flow Cytometry"

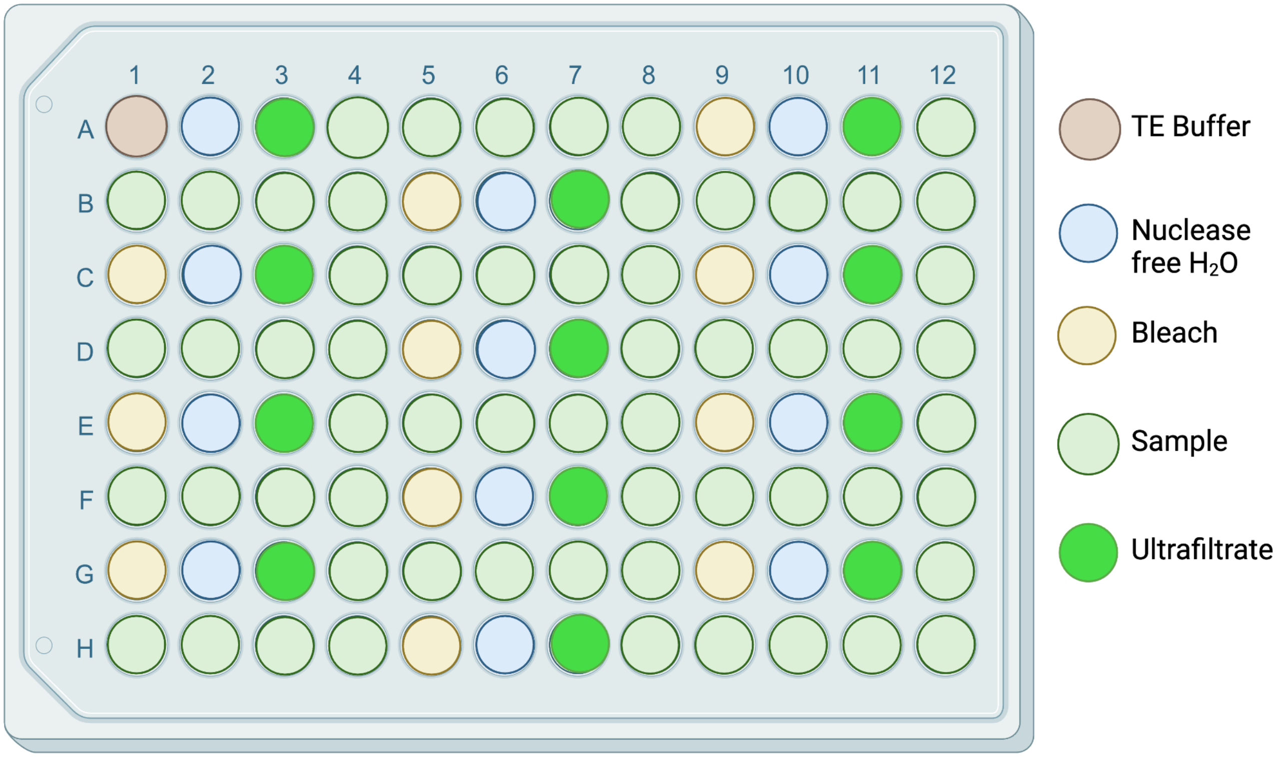


**Supplemental Figure 1.** Visual representation of the well plate plan. Well A1 is filled with TE buffer as a blank to determine the machine's base noise level. Twelve wells are filled with nuclease free water to rinse the cytometer and manage noise levels of the machine. Twelve wells are filled with ultrafiltrate that was treated the same as the sample to determine the base noise level of the sample. Eleven wells are filled with a 10% bleach solution to cleanse the machine throughout the run. Sixty wells are filled with the diluted sample. Throughout the well plate, a set of three wells containing 10% bleach, nuclease free water, and ultrafiltrate occur for every five sample wells. This ensures that an overestimation of VLPs are not being counted and the noise levels can be constantly managed.

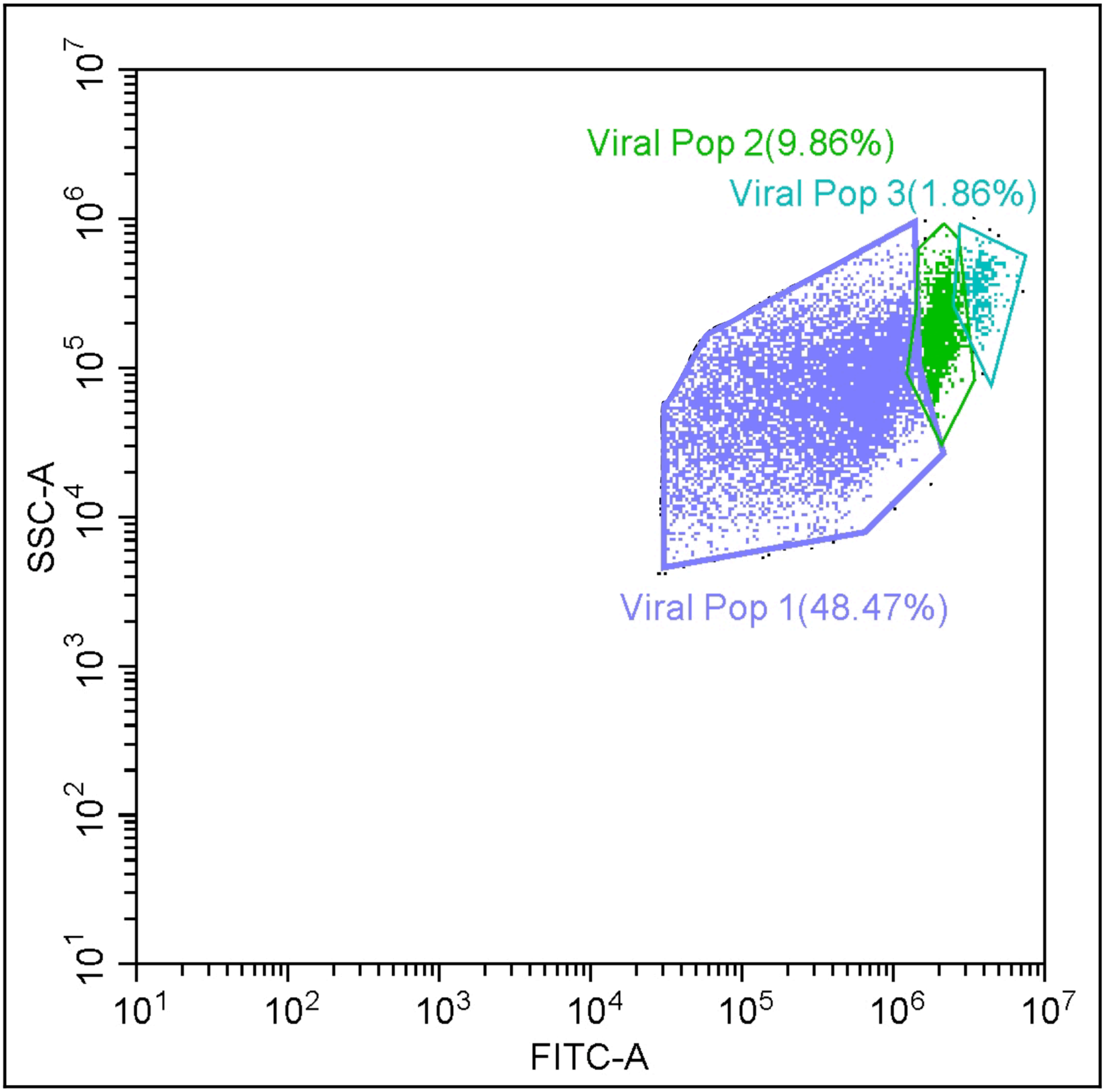


**Supplemental Figure 2.** Three distinct viral populations in a sample after treatment with chloroform and benzonase. Populations are identified and gated by looking for clusters of events (shown as dots) in the region of interest. Here, viral populations should have a high fluorescent intensity (FITC-A, X axis) and medium-high side scatter (SSC-A, Y axis). In this plot, only the gates of interest are shown.

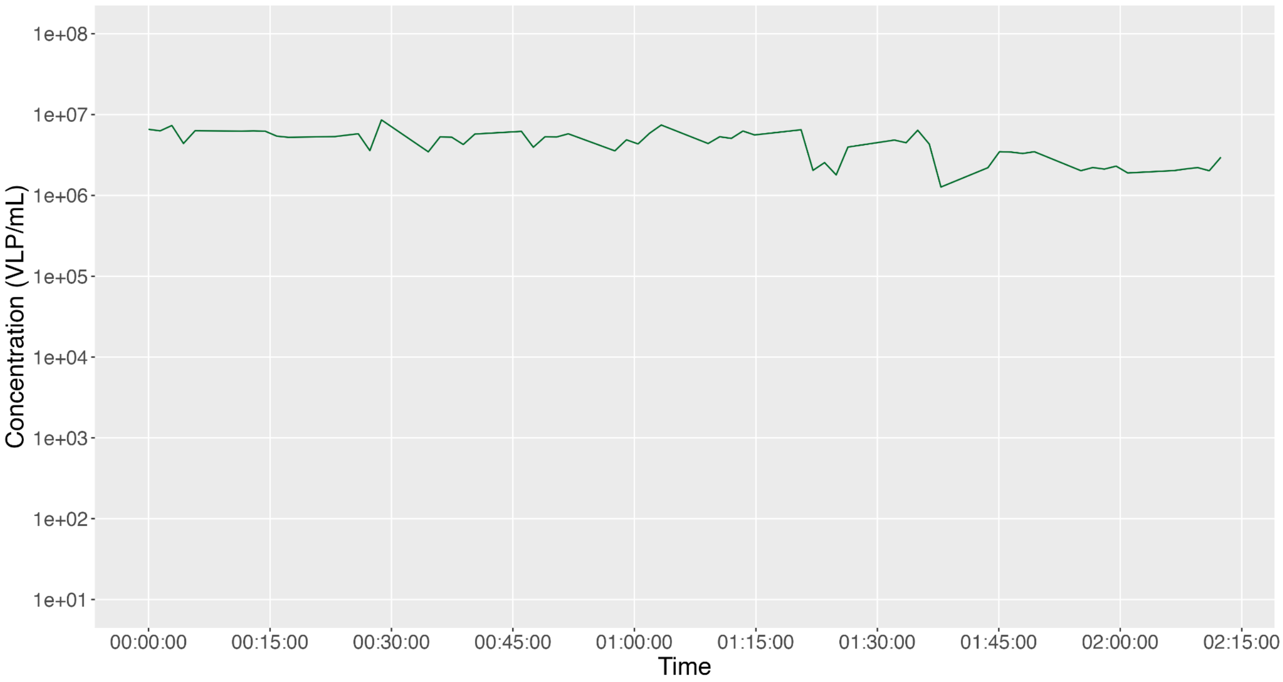


**Supplemental Figure 3.** Untreated WBS sample's VLPs $mL^{-1}$ over the course of the well plate's run on the flow cytometer. Time points are plotted for each well in reference to the time elapsed between that well and the first sample well, when T=0 hours, 0 minutes, 0 seconds (00:00:00). Two points of significant degradation can be seen between 1:15:00 and 1:30:00 and between 1:30:00 and 1:45:00.

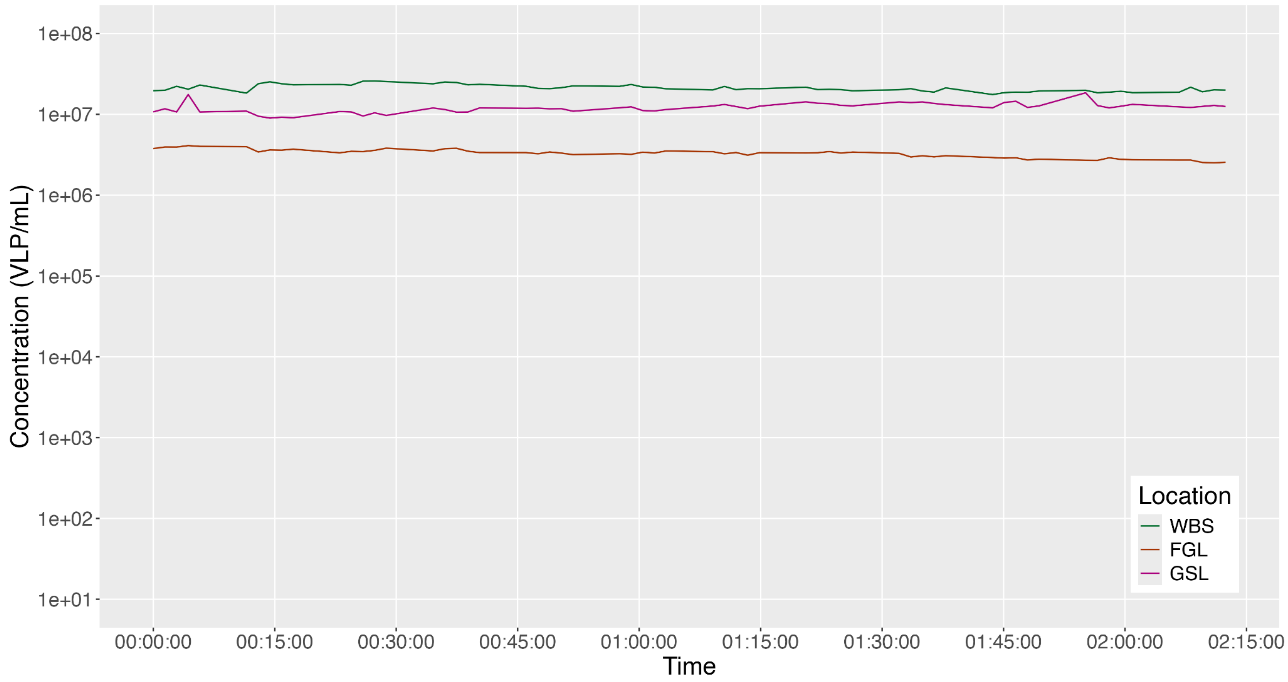


**Supplemental Figure 4.** Each sample location's VLP mL$^{-1}$ over the course of the well plate's run on the flow cytometer. Samples were treated with chloroform and benzonase. FGL is shown in orange (bottom). GSL is shown in pink (middle). WBS is shown in green (top). Time points are plotted for each well in reference to the time elapsed between that well and the first sample well, when T=0 hours, 0 minutes, 0 seconds (00:00:00).

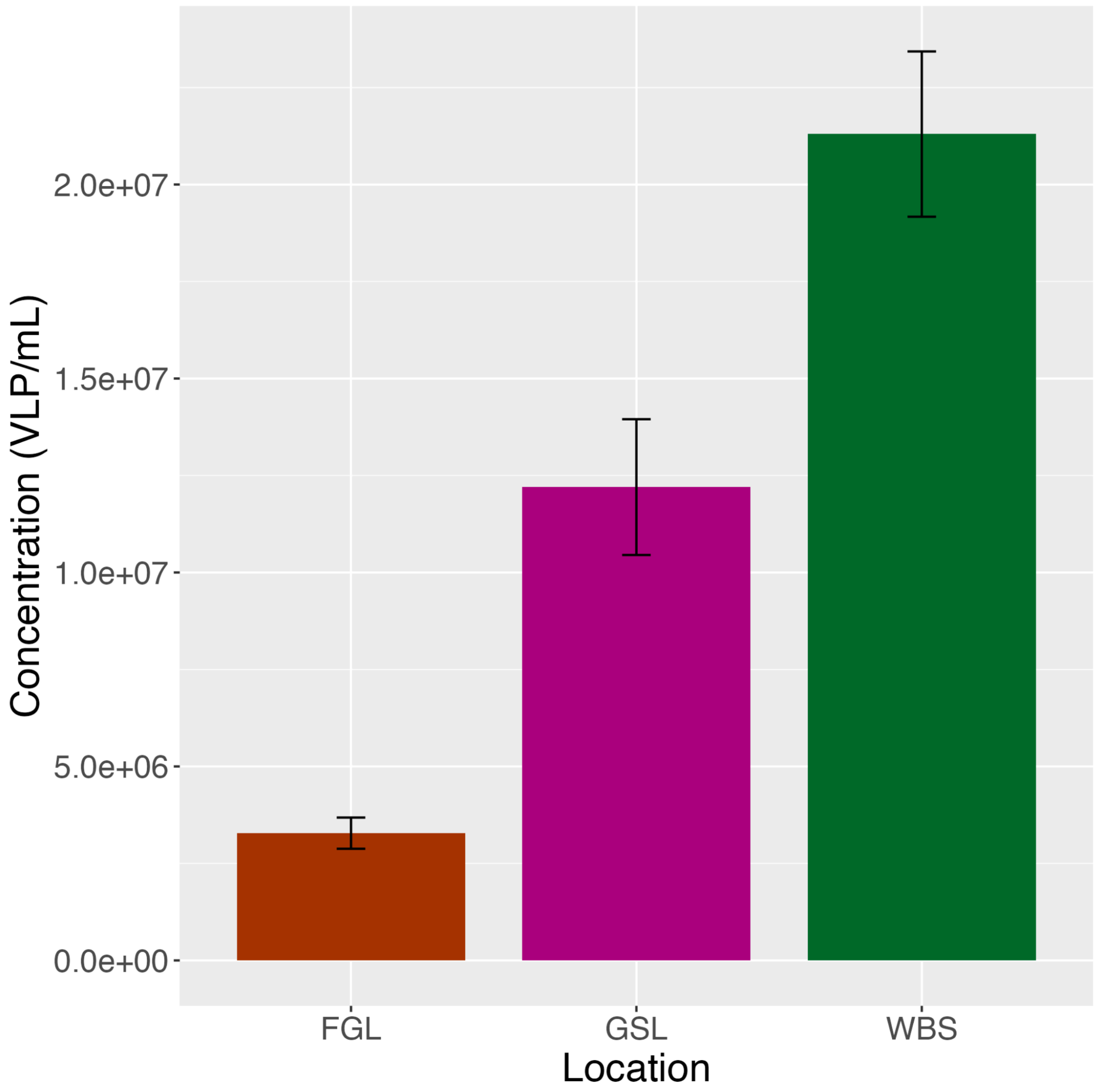


**Supplemental Figure 5.** Average VLP $mL^{-1}$ at each sample location after chloroform and benzonase treatment. Error bars represent standard deviation values.

**Supplemental Table 1.** Results from the statistical analysis of the untreated FGL experiment to identify degradation. Each row was analyzed and compared to the first row to find significant differences. No significant degradation was detected.

**Supplemental Table 2.** Results from the statistical analysis of the untreated GSL experiment to identify degradation. Each row was analyzed and compared to the first row to find significant differences. No significant degradation was detected.

**Supplemental Table 3.** Results from the statistical analysis of the untreated WBS experiment to identify degradation. Each row was analyzed and compared to the first row to find significant differences. Significant degradation was detected in rows F, G, and H. A dynamic sliding window of three wells throughout rows F, G, and H were analyzed to find the exact point of degradation. Comparisons were made to the first set of three wells in the experiment. Degradation at the beginning of rows F and G were detected and sustained for four wells and eleven wells, respectively.

**Supplemental Table 4.** Average VLP concentrations at each location for each experiment type. Experiments ending in "-C" are chloroform treated samples. Experiments beginning with "B-" are benzonase treated samples. Averages and standard deviations are represented as VLPs $mL^{-1}$. Comparisons of averages obtained using EFM are also presented.
